# The genome analysis of *Tripterygium wilfordii* reveals TwCYP712K1 and *TwCYP712K2* responsible for oxidation of friedelin in celastrol biosynthesis pathway

**DOI:** 10.1101/2020.06.29.176958

**Authors:** Tianlin Pei, Mengxiao Yan, Yu Kong, Jie Liu, Mengying Cui, Yumin Fang, Binjie Ge, Jun Yang, Qing Zhao

## Abstract

*Tripterygium wilfordii* is a Traditional Chinese Medicine (TCM) from family Celastraceae and celastrol is one of the strongest active ingredients belonging to friedelane-type pentacyclic triterpenoid, which has a large clinical application value of anti-tumor, immunosuppression, and obesity treatment. The first committed biosynthesis step of celastrol is the cyclization of 2, 3-oxidosqualene to friedelin, catalyzed by the oxidosqualene cyclase, while the rest of this pathway is still unclear. In this study, we reported a reference genome assembly of *T. wilfordii* with high-quality annotation by using a hybrid sequencing strategy (Nanopore, Bionano, Illumina HiSeq, and Pacbio), which obtained a 340.12 Mb total size and contig N50 reached 3.09 Mb. In addition, we successfully anchored 91.02% sequences into 23 pseudochromosomes using Hi-C technology and the super-scaffold N50 reached 13.03 Mb. Based on integration genome, transcriptom and metabolite analyses, as well as *in vivo* and *in vitro* enzyme assays, two CYP450 genes, *TwCYP712K1* and *TwCYP712K2* have been proven for C-29 position oxidation of friedelin to produce polpunonic acid, which clarifies the second biosynthesis step of celastrol. Syntenic analysis revealed that *TwCYP712K*1 and *TwCYP712K2* derived from the common ancestor. These results have provided insight into illustrating pathways for both celastrol and other bioactive compounds found in this plant.

## INTRODUCTION

*Tripterygium wilfordii* Hook. f. is a perennial twining shrub that belongs to the Celastraceae family and is known as “Lei gong teng” (means Thunder God Vine) in China. It is indigenous to southeast of China, Korean Peninsula, Japan and has been cultivated worldwide as a medicinal plant nowadays (Helmstaedter, 2013; Tian et al., 2019). In Chinese, Lei Gong means Thunder God, the name probably indicates its strong biological activity along with high toxicity that reminds doctors use it carefully (Gao et al., 2012). For centuries, Chinese people have used the extraction of *T. wilfordii* bark as a kind of pesticide because of its toxicity, as recoded in Illustrated Catalogue of Plants, firstly published in 1848 (Wu, 1957). The medicinal importance of *T. wilfordii* was recognized in 1960s, for its root successfully relieved symptoms of leprosy patients in Gutian County, Fujian province of China (Gutian-leprosy-hospital, 1972). This event spurred interest of researchers in various areas. *T. wilfordii* was then proven to be effective in treatment of autoimmune diseases such as rheumatoid arthritis and systemic psoriasis (Bao and Dai, 2011; Han et al., 2012). In recent decades, many studies have shown that the extraction of *T. wilfordii* has anti-cancer, anti-diabetic and anti-inflammatory effects (Ding et al., 2015; Gao et al., 2010; Yuan et al., 2017).

The pharmacological activities of *T. wilfordii* were mainly attributed to the various compounds accumulate in its root, such as alkaloid, diterpenoids and triterpenoids (Duan et al., 2000; Lv et al., 2019). Celastrol belongs to friedelane-type triterpenoid mainly found in root bark of *T. wilfordii* (Lange et al., 2017). It has been shown to be effective in the treatment of inflammatory and autoimmune diseases (Ma et al., 2007), anti-tumors (Pang et al., 2019), and as a possible treatment for Alzheimer’s disease (Allison et al., 2001). Besides, celastrol has found to be a leptin sensitizer and a powerful compound for the treatment of obesity (Liu et al., 2015; Ma et al., 2015). Despite the importance of natural products found in *T. wilfordii* and their growing demands, the traditional production is becoming unsustainable, for its slow growing rate and low accumulation of celastrol (Lin et al., 2009). This situation strongly argues novel producing methods such as synthetic biology. Genome sequencing will offer us a reference to mine the genes responsible for the pathways of the bioactive compounds.

Celastrol is pentacyclic triterpenoid synthesized from 2, 3-oxidosqualene, which is the common biosynthetic precursor of triterpenoid derived from the cytosolic mevalonate (MVA) and plastid 2-C-methyl-D-erythritol-4-phosphate (MEP) pathway (Miettinen et al., 2017a; Thimmappa et al., 2014). Two oxidosqualene cyclase (OSC), TwOSC1 and TwOSC3, are identified as key enzymes in cyclization of 2, 3-oxidosqualene to form friedelin, which is the first committed step in celastrol formation (Zhou et al., 2019). The subsequent step in this pathway is consider to be the hydroxylation of C-29 position of friedelin to produce 29-hydroxy-friedelin-3-one and turn into polpunonic acid by carboxylation, which is then undergone a serious of oxidation and rearrangements to produce celastrol (Zhou et al., 2019).

Cytochrome P450 monooxygenases (CYPs) are the largest enzyme family in plants and devote to execute specialized metabolism by backbone modification of compounds including oxidation of triterpenoid scaffold, which is important for the diversification and functionalization of the natural products (Bathe and Tissier, 2019; Ghosh, 2017). There are 127 CYP families from plants are identified and they are categorized into 11 clans of which include single or multiple sub-clades on a phylogenetic tree (Nelson and Werck, 2011). So far, many CYP families are reported to participate in pentacyclic triterpenoid structural modifications, including CYP51, CYP71, CYP72, CYP87, CYP88, CYP93, and CYP716 (Ghosh, 2017; Miettinen et al., 2017b; Ribeiro et al., 2020).

Here, we report the reference genome assembly of *T. wilfordii* by using a combined sequencing strategy. After integrating genome, transcriptom and metabolite analyses, a number of CYP genes highly related to celastrol biosynthesis were identified. TwCYP712K1 and TwCYP712K2 were then functionally characterized as oxidization of the C-29 position of friedelin and production of polpunonic acid by using *in vivo* yeast and *in vitro* enzyme assays. These results pave an important strategy to elucidate pathways not only for celastrol, but also for other bioactive compounds found in this plant.

## RESULTS

### Genome Sequencing, Assembly, and Annotation

We obtained 77.86 Gb of Nanopore reads, amounting to 207.16× coverage of the estimated genome size (375.84 Mb) of genome survey (Supplementary Figure 1 and Supplementary Table 1). Draft genome was assembled to obtain primary contigs, with 340.12 Mb total size and 3.09 Mb contig N50 (Supplementary Table 2). GC content of the genome was 37.19%, with N comprising 0.00% (Supplemental Table 3). Variant calling showed a heterozygosity rate of 0.25% (Supplemental Table 4). Benchmarking Universal Single-Copy Orthologs (BUSCO) analysis showed 95.2% of complete single-copy genes (Supplemental Table 5). Unsurprisingly, Core Eukaryotic Genes Mapping Approach (CEGMA) analysis gained 238 core eukaryotic genes (95.97%, Supplemental Table 6). Both results indicated that the presented genome is relative complete. The short reads obtained from Illumina sequencing of genome survey were aligned to the genome (Supplemental Table 2), which exhibited a high consistency of 95.31% mapping rate and 93.99% coverage (Supplemental Table 7). For assembly improvement, 60.80 Gb (161.77× of estimated genome size) reads from BioNano sequencing were obtained and integrated with draft genome to construct scaffolds, which updated the genome of *T. wilfordii* from a total contig length of 340.12 Mb, a contig N50 of 3.09 Mb, to a total scaffold length of 342.59 Mb and a scaffold N50 of 5.43 Mb (Table 1 and Supplemental Table 2). Furthermore, we anchored 91.02% of the assembly into 23 pseudochromosomes by using Hi-C technology (Supplemental Table 8). All the super-scaffold could be placed in one of twenty three groups (Supplemental Figure 2). The super-scaffold N50 reached 13.03 Mb, with the longest super-scaffold being 17.75 Mb (Supplemental Table 8 and 9).

**Table 1.** Summary of *de novo* genome assembly and annotation of *T. wilfordii*.

|  | Number | Index |
| --- | --- | --- |
| **Genome assembly** |  |  |
| Total contigs | 553 | 340.12 Mb |
| Contig N50 | 34 | 3.09 Mb |
| Total scaffolds | 470 | 342.59 Mb |
| Scaffold N50 | 25 | 5.43 Mb |
| GC content |  | 37.19% |
| Heterozygosity rate |  | 0.25% |
| Pseudochromosomes | 23 | 311.85 Mb |
| Super-scaffold N50 |  | 13.03 Mb |
| **Genome annotation** |  |  |
| Repetitive sequences | 44.31% | 151.81 Mb |
| Noncoding RNAs | 4770 | 0.82 Mb |
| Structure genes | 31593 | 100.48 Mb |

For genome annotation and gene expression profile analyses, roots (R), stems (S), young leaves (YL), mature leaves (L), flower buds (FB) and flowers (F) of *T. wilfordii* plants were collected prior to RNA-seq, using Illumina platform. Furthermore, we mixed the RNA samples of different tissues and sequenced using PacBio platform to obtain the full-length transcriptome sequences (Supplemental Table 2). A combined strategy containing *de novo* prediction, homologous prediction, RNA-seq and PacBio read alignment was used to construct gene structure for the *T. wilfordii* genome. The final set of annotated genes amounted to 31593, with an average length of 3180 bp and an average coding sequence length of 1182 bp (Supplemental Table 10). A total of 27301 genes (86.41%) were supported by RNA-seq data, and 23229 genes (73.53%) were supported by all used methods, and these genes were annotated with high confidence. Gene function annotation was performed by blasting the protein sequences of predicted genes against the public database, including NR, SwissProt, KEGG, GO, Pfam and InterPro. A total of 30535 (96.70%) gene products could be function predicted, and 22491 sequences could be annotated by at least one of the databases (NR, SwissProt, InterPro and KEGG) (Supplemental Table 11).

Repeat sequences annotation showed that the genome of *T. wilfordii* contained 44.31% repetitive sequences. Among these sequences, tandem repeat (small satellites and microsatellites) and interspersed repeats accounted for 0.95% and 43.36%, respectively. Long terminal repeats (LTRs) of retroelements were the most abundant interspersed repeat, occupying 36.74% of the genome, followed by DNA transposable elements at 1.68% (Supplemental Table 12). Non-coding RNA annotation revealed that the *T. wilfordii* genome possessed a total of 355 microRNAs (miRNAs), 797 transfer RNAs (tRNAs), 827 ribosomal RNAs (rRNAs), and 982 small nuclear RNAs (snRNAs), respectively (Supplemental Table 13). The integrated distributions of the genes, repeats, non-coding RNA densities, and all detected segmental duplications are shown in Figure 1.

**Figure 1.**
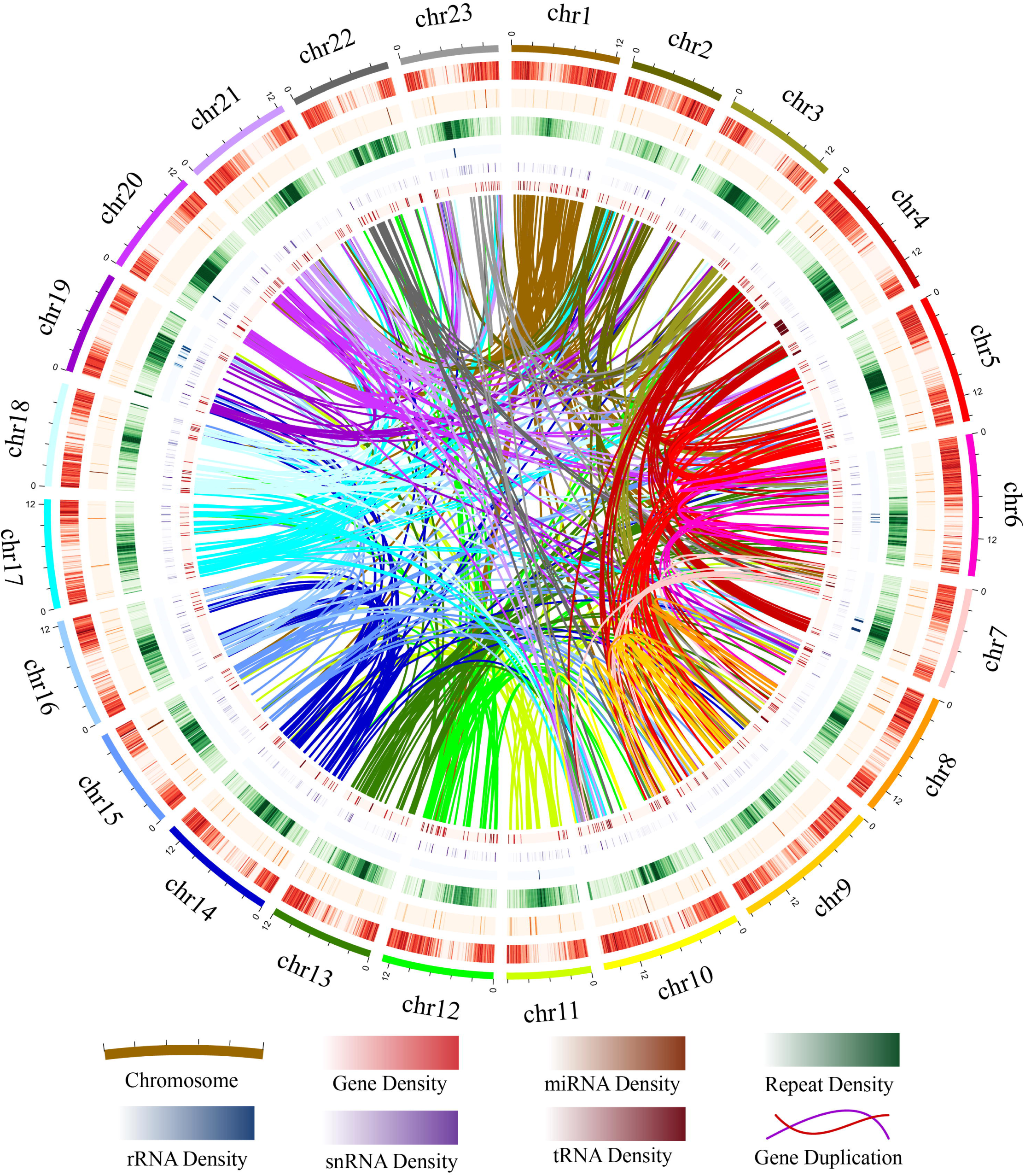
Landscape of *T. wilfordii* genome assembly. Circles from outer to inner represent: pseudochromosomes, density of genes, miRNAs, repeats, rRNAs, snRNAs, tRNAs, and duplicated gene links within genome, respectively. Scale shows chromosomes in a 500 kb window; gene density in a 100 kb window (0–100%, which means the percentage of gene density indicated by the color gradient starts from 0 and goes to 100% of 100 kb DNA); miRNA density in a 100 kb window (0–1%); repeat density in a 100 kb window (0–100%); rRNA density in a 100 kb window (0–18%); snRNA density in a 100 kb window (0–1.3%); tRNA density in a 100 kb window (0–1.8%); detected gene duplication links (570).

### Comparative Genomic Analysis

To identify evolutionary characteristics and gene families, the genome of *T. wilfordii* was compared with 11 published genomes of nine eudicots species (*Arabidopsis thaliana, Citrus sinensis, Vitis vinifera, Glycine max, Medicago truncatula, Glycyrrhiza uralensis, Cucumis sativus, Populus trichocarpa* and *Ricinus communis*) and a monocot species (*Oryza sativa*), plus *Amborella trichopoda*, one of the basal group of angiosperms, was selected as an outgroup. Based on analysis of gene family clustering, we identified 29189 gene families, of which 7296 were shared by all 12 species, and 485 of these shared families were single-copy gene families (Supplemental Figure 3). We compared the gene family numbers among *T. wilfordii* and four fabids species. As shown in Figure 2A, 10722 gene families were shared by *G. max, C. sativus, G. uralensis*, and *M. truncatula*, and 1086 gene families were specific to *T. wilfordii*. When comparing with their most recent common ancestor (MRCA) of the 12 plant species, 15 gene families with 152 genes were expanded and 42 gene families with 54 genes were contracted in *T. wilfordii* (Figure 2B). KEGG studies showed that the expanded genes enriched in pathways related to “ubiquinone and other terpenoid-quinone biosynthesis” and “steroid hormone biosynthesis”, suggesting gene family expansion contributed to specialized metabolite biosynthesis in *T. wilfordii*. Phylogenetic tree was constructed based on the superalignment matrix of 485 single-copy orthologous genes from the 12 species. The branching order showed that *A. thaliana* (Brassicales) and *C. sinensis* (Sapindales) were sister to *P. trichocarpa, R. communis* (Malpighiales) and *T. wilfordii* (Celastrales), which diverged approximately 109.1 million years ago (Mya), followed by divergence of *T. wilfordii* and species from Malpighiales about 102.4 Mya (Figure 2B and Supplemental Figure 4). These results were consistent with a previously proposed phylogenetic order which Celastrales and Malpighiales formed to be sister to each other (Zhao et al., 2016a).

**Figure 2.**
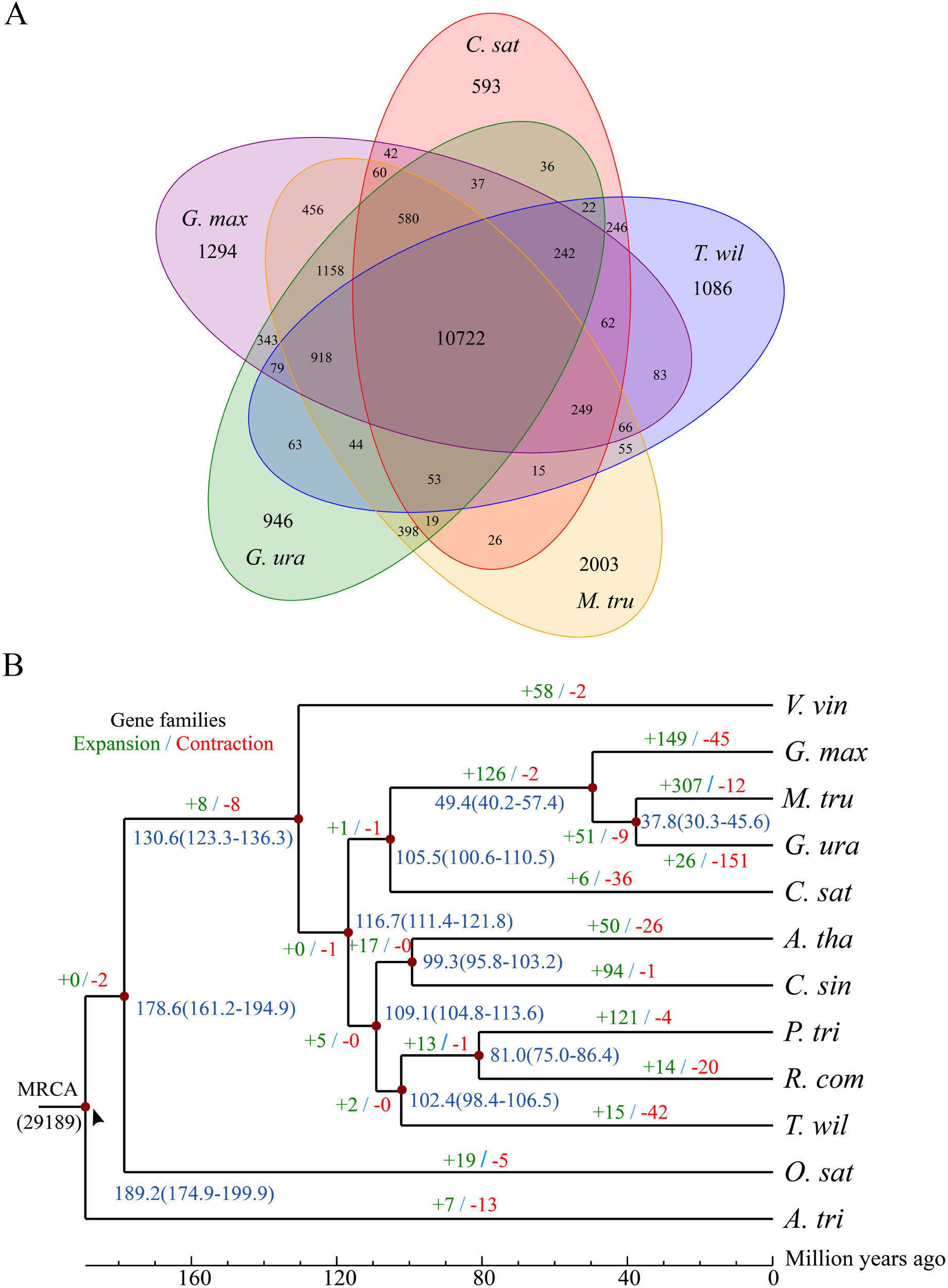
Comparative genomic analysis. (A) Venn diagram of common and unique gene families in *T. wilfordii* with that in other species. *T. wil* = *Tripterygium wilfordii*; *G. max* = *Glycine max*; *M. tru* = *Medicago truncatula*; *G. ura*= *Glycyrrhiza uralensis*; *C. sat* = *Cucumis sativus*. (B) Phylogenetic analysis, divergence time estimation, and gene family expansions and contractions. Divergence times (Mya) are indicated by the blue numbers, and numbers in brackets represent confidence interval. Gene family expansions and contractions are indicated by green and red numbers, respectively. *T. wil* = *Tripterygium wilfordii*; *A. tha* = *Arabidopsis thaliana*; *O. sat* = *Oryza sativa*; *R. com* = *Ricinus communis*; *P. tri* = *Populus trichocarpa*; *G. max* = *Glycine max*; *M. tru* = *Medicago truncatula*; *G. ura*= *Glycyrrhiza uralensis*; *V. vin* = *Vitis vinifera*; *C. sin* = *Citrus sinensis*; *C. sat* = *Cucumis sativus*; *A. tri* = *Amborella trichopoda*.

### Genome-wide identification and analysis of CYP450 candidates involved in celastrol biosynthesis

Based on the HMMER analysis, we totally extracted 213 full-length ORFs of CYP genes from *T. wilfordii* genome and annotated by phylogenetic analysis with CYPs from *A. thaliana*. We identified 35 CYPs related to triterpenoid oxidases belonging to different subfamily, including CYP716, CYP72, CYP71, CYP93, CYP705A and CYP81 which were reported functionally associating with diverse triterpenoid structural modifications (Figure 3) (Ghosh, 2017).

**Figure 3.**
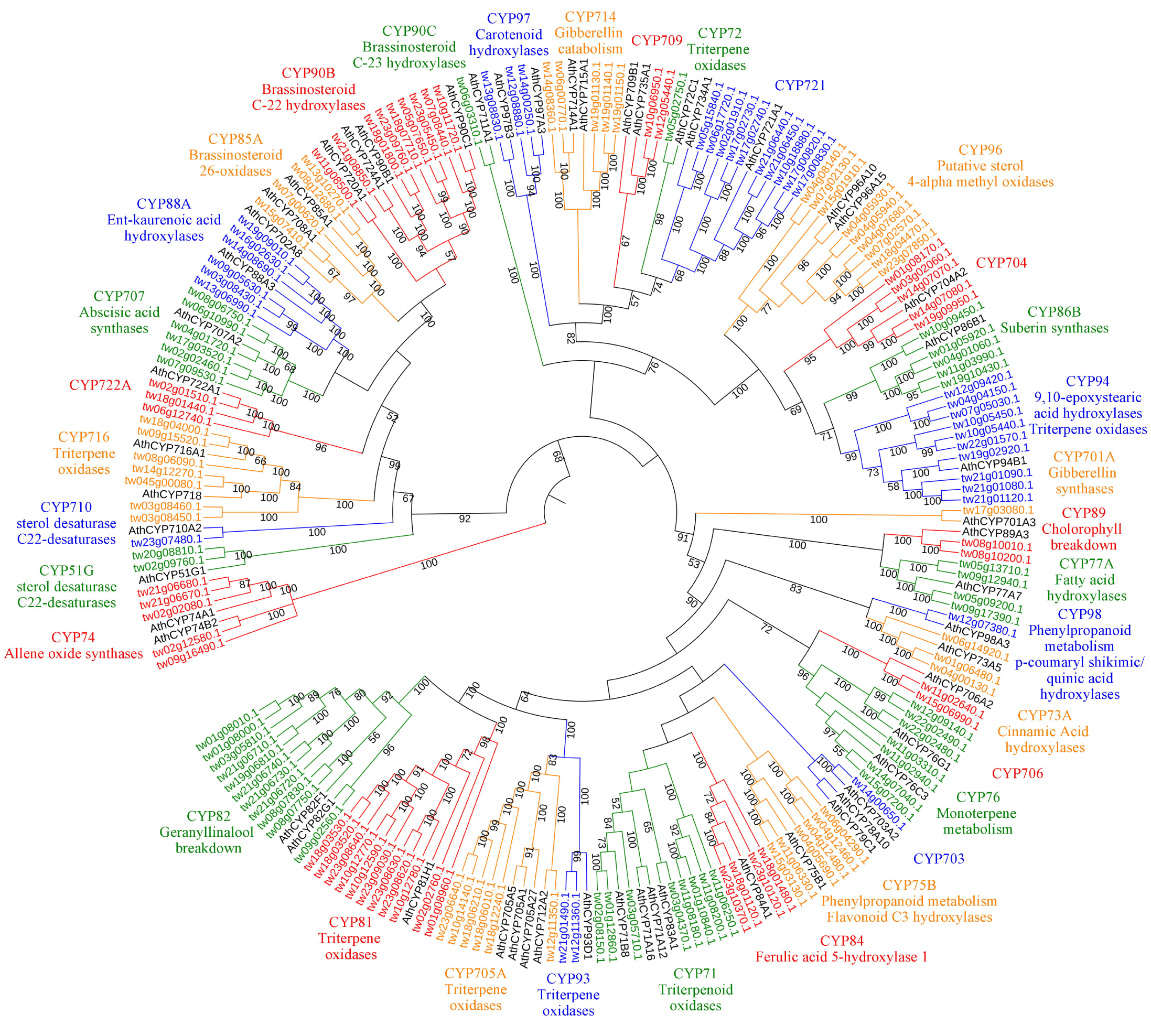
Phylogenetic tree of *T. wilfordii* CYPs. Diverse functions of CYPs are annotated by CYPs from *A. thaliana*. Phylogenetic tree was built using the Neighbor-Joining methods with a bootstrap test (n=1000 replications). Numbers on the branches represent bootstrap support value.

On the other hand, we investigated the expression patterns of the identified 213 *CYPs* with *TwOSC1* and *TwOSC3*, which were the two committed enzymes involved in precursor of celastrol biosynthesis in *T. wilfordii* (Zhou et al., 2019). Based on the RNA-Seq data of various tissues, we obtained 20 coexpression profiles of which only profile #3 and #13 showed significance (*P*-value<0.05) (Supplemental Figure 5). Profile #3 contained *TwOSC3* and 45 *CYPs* showed similar expression pattern, while profile #13 included *TwOSC1* and 51 *CYPs* had coexpression trends (Supplemental Figure 6), which suggested that these *CYPs* potentially involved in the biosynthesis of celastrol.

To narrow down the candidate genes, we compared the 35 CYPs identified by phylogenetic analysis and the genes showed coexpression patterns with *TwOSC1* and *TwOSC3*, respectively. As shown in Figure 4A, there were nine and seven triterpenoid biosynthesis-related *CYPs* showed coexpression patterns with *TwOSC1* and *TwOSC3*, respectively. However, no *CYPs* were crossover between *TwOSC1* group and *TwOSC3* group, suggesting that two independent pathway of friedelane-type triterpenoid biosynthesis mediated by *TwOSC1* and *TwOSC3*, respectively. Based on the tissue expression profiles, the 16 *CYPs* were clustered into two clades with *TwOSC1* and *TwOSC3* separately, which *TwOSC3* was a root-specific expression gene, while *TwOSC1* highly expressed in leaf and other aerial parts (Figure 4B).

We calculated the gene to gene and gene to metabolite Pearson’s correlation coefficients (r) by using tissue expression profiles of 16 outstand *CYPs* mentioned above, and three other known genes related to celastrol biosynthesis (*TwHMGR1, TwFPS1* and *TwDXR*) (Wang et al., 2018; Zhang et al., 2018; Zhao et al., 2017), known intermediate product and celastrol concentrations (Zhou et al., 2019). As shown in Figure 4C, seven *CYPs* were positively correlated to celastrol biosynthesis-related genes with high Pearson’s r and significant *P*-value. Besides, these *CYPs* also highly correlated with 29-hydroxy-friedelan-3-one, polpunonic acid, wilforic acid A and celastrol contents which were all specific accumulated in the roots of *T. wilfordii* (Supplemental Figure 7).

**Figure 4.**
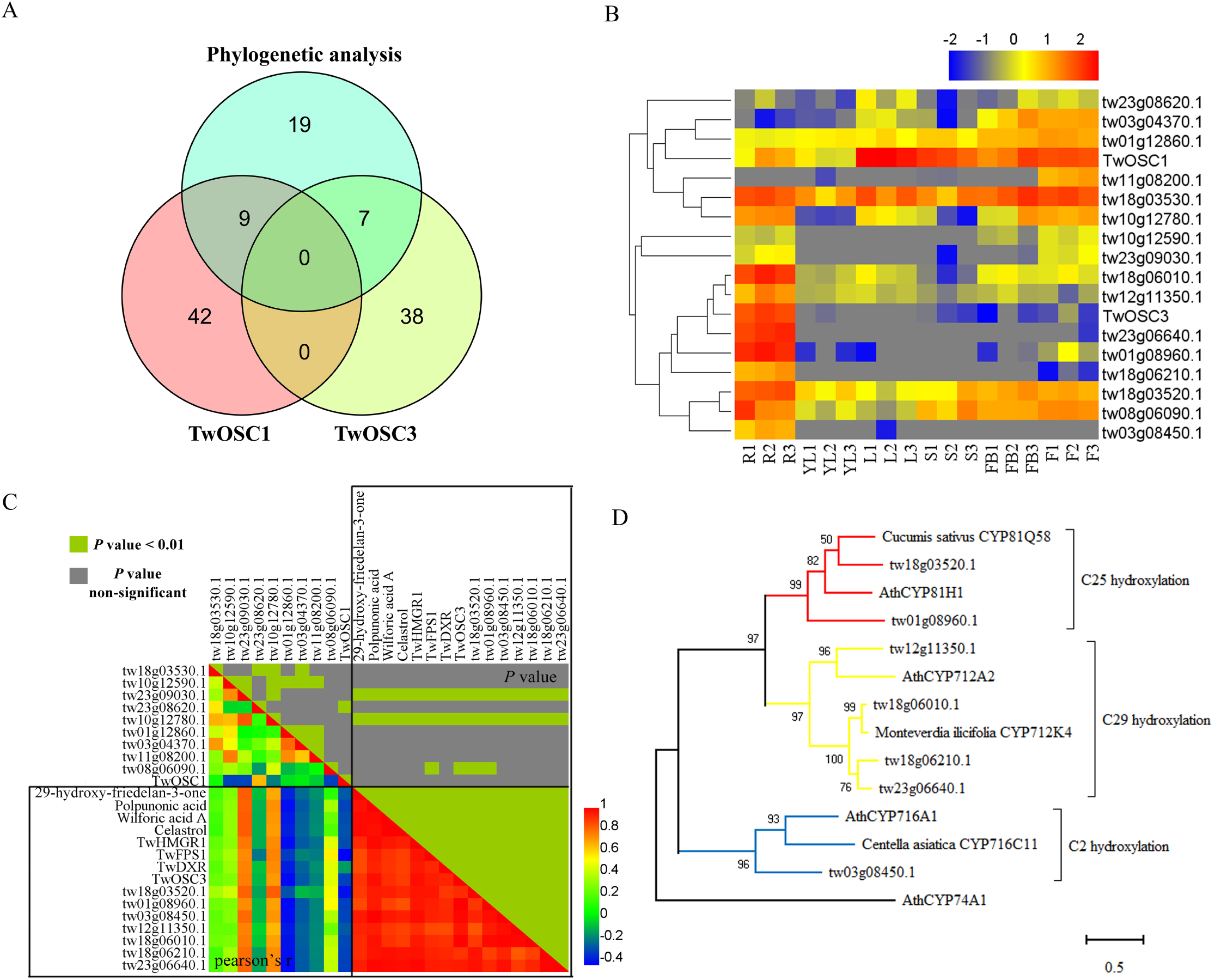
Identification of polpunonic acid producing *CYPs* via integration analysis. (A) Venn diagram of identified CYPs by phylogenetic analysis versus coexpression patterns. (B) Tissue specific expression profiles and clustering of *CYPs* with *TwOSC1* and *TwOSC3*. Gradient bar represents the expression levels from high (red) to low (blue) with log2 normalization, and color gray represents the empty value. R, root; YL, young leaf; L, leaf; S, stem; FB, flower bud; F, flower; numbers 1-3 represents three biological replicates, respectively. (C) Matrix of Pearson’s correlation coefficient and corresponding *P*-value of compounds, biosynthesis-related genes and *CYPs* candidates. The lower triangle matrix represents Pearson’s correlation coefficient, and the upper triangle matrix presents *P*-values test (green indicates *P*-values < 0.01, and gray indicates non-significant correlation). Gradient bar represents the correlation coefficient of positive or negative correlation from high to low. The black boxes enclose the highly correlated genes or compounds. (D) Phylogenetic analysis of putative CYPs in celastrol biosynthesis and the CYPs known to catalyze structural modifications on triterpenoid scaffolds Phylogenetic tree was built using the Maximum-likelihood methods with a bootstrap test (n=1000 replications). Allene oxide synthases AtCYP74A1 was set as an outgroup.

Phylogenetic analysis placed these seven *CYPs* and *CYPs* from other species into three clades which represented different function in triterpenoid structural modifications (Figure 4D). Two CYPs (tw18g03520.1 and tw01g08960.1) were clustered with CYP81Q58 from *C. sativus* which catalyzed the C-25 position hydroxylation of cucurbitacins (Shang et al., 2014), four CYPs (tw12g11350.1, tw18g06010.1, tw18g06210.1 and tw23g06640.1) were clustered with CYP712K4 from *Monteverdia ilicifolia* which catalyzed the C-29 position oxidation using friedelin as a substrate (Bicalho et al., 2019), and tw03g08450.1 was clustered with CYP716C11 from *Centella asiatica* which hydroxylated C-2 position of oleanolic acid and ursolic acid (Miettinen et al., 2017b).

Since we were interested in looking for C-29 position oxidase that could catalyze friedelin into polpunonic acid, we finally chose 3 *CYPs* as candidates to do function validation, including tw18g06010.1, tw18g06210.1 and tw23g06640.1 (named *TwCYP712K1, TwCYP712K2* and *TwCYP712K3* hereafter) according to the closer relationship with CYP712K4 from *M. ilicifolia* related to C-29 hydroxylation (Bicalho et al., 2019).

### Heterologous expression and characterization of putative CYPs

We successfully cloned the full-length ORFs of *TwCYP712K1, TwCYP712K2* and *TwCYP712K3*, and then separately expressed in yeast which was fed with friedelin or 29-hydroxy-friedelan-3-one, respectively. However, we could not detect any new peak from the yeast strains expressing the enzymes supplemented with friedelin compared to the empty vector (EV) control (data not shown), which probably because the hydrophobic substrate was unable to transport in to yeast cell. When fed with 29-hydroxy-friedelan-3-one, both TwCYP712K1 and TwCYP712K2 converted the substrate to polpunonic acid, while TwCYP712K3 did not show such activity in this assay (Figure 5A and 5D). To further explore the enzyme activities, we extracted the microsomes from yeast cells and incubated the proteins with friedelin for 12 h. As shown in Figure 5B, polpunonic acid was detected in both TwCYP712K1 and TwCYP712K2 incubation, which supported our previous speculation that friedelin supplemented in media may be unable to transport in to yeast cell. In addition, TwCYP712K1 and TwCYP712K2 converted 29-hydroxy-friedelan-3-one to polpunonic acid *in vitro* (Figure 5C), consisting with previous yeast *in vivo* assays. These results demonstrated that both TwCYP712K1 and TwCYP712K2 can catalyze two-step oxidation of friedelin to generate 29-hydroxy-friedelan-3-one and polpunonic acid in celastrol biosynthesis (Figure 5E).

**Figure 5.**
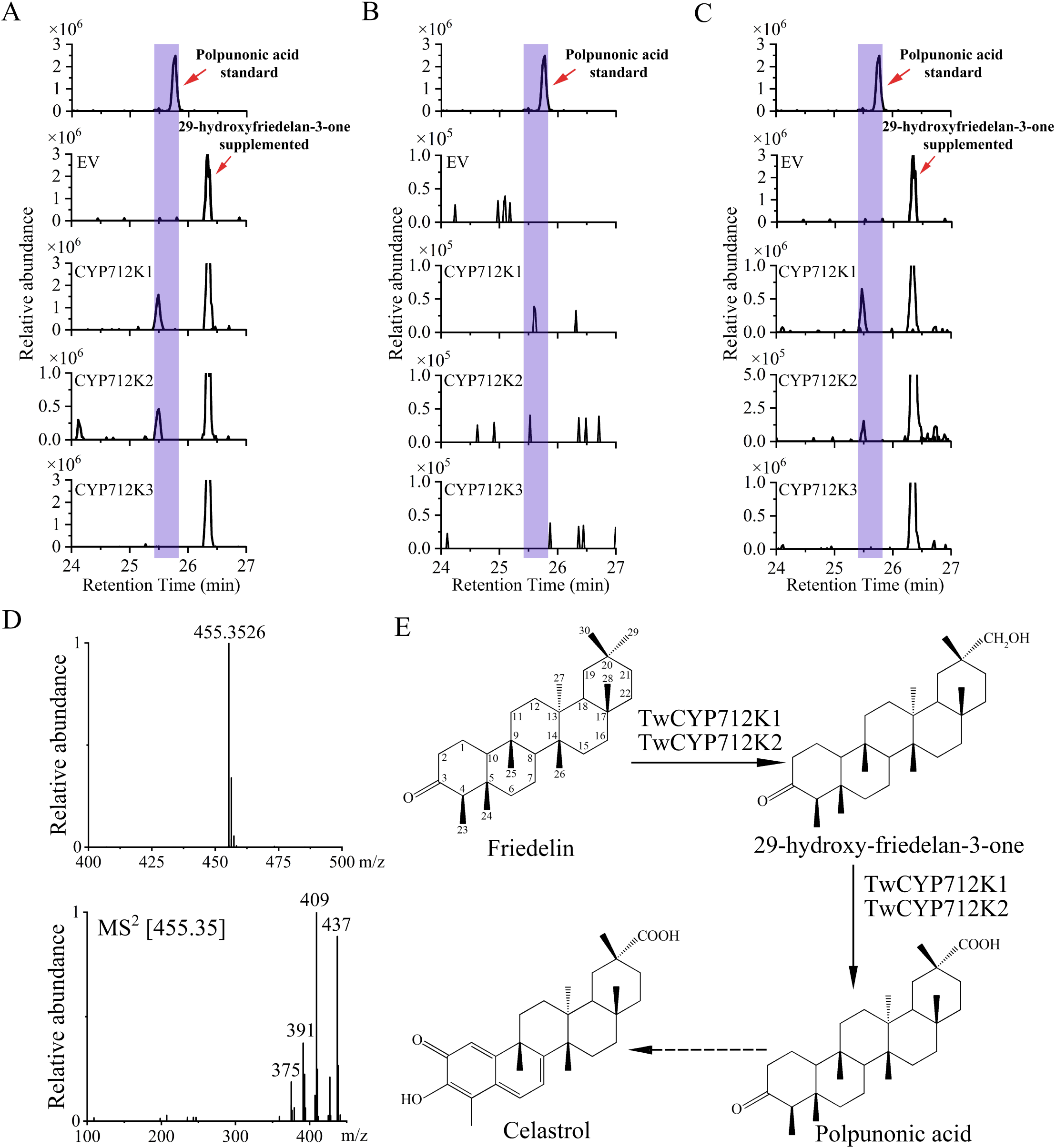
Characterization of candidate CYPs for polpunonic acid biosynthesis. (A) LC-MS analysis of yeast samples fed with 29-hydroxy-friedelan-3-one as substrate in vivo. Top, polpunonic acid standard; EV, empty vector; TwCYP712K1-TwCYP712K3, yeast expressing the corresponding proteins. New peaks with the same retention time as polpunonic acid were highlight. (B) LC-MS analysis of microsomes samples incubated with friedelin as substrate in vitro. Top, polpunonic acid standard; EV, empty vector; TwCYP712K1-TwCYP712K3, corresponding microsomes extracted from yeast cells. New peaks with the same retention time as polpunonic acid were highlight. (C) LC-MS analysis of microsomes samples incubated with 29-hydroxy-friedelan-3-one as substrate in vitro. Top, polpunonic acid standard; EV, empty vector; TwCYP712K1-TwCYP712K3, corresponding microsomes extracted from yeast cells. New peaks with the same retention time as polpunonic acid were highlight. (D) Accurate masses of polpunonic acid (top) and its MS^2^ fragmentation pattern (bottom). (E) Two-step oxidation of friedelin catalyzed by TwCYP712K1 and TwCYP712K2. The dashed arrow indicates multiple catalyzed steps which were unidentified.

### Evolution analyses of *TwCYP712K1* and *TwCYP712K2*

Genome analysis showed that *TwCYP712K1* (tw1806010.1) and *TwCYP712K2* (tw18g06210.1) located in pseudochromosome 18 within an approximately 200 kb region, which inspired us to examine the evolutionary relationship between the specific CYPs. We proposed that must be a gene duplication event during evolution. However, amino acid alignment indicated that only 70.57% identity between TwCYP712K1 and TwCYP712K2 (Supplementary Figure 8), suggesting that these two genes specialized long time ago. Syntenic analysis showed that TwCYP712K1 has corresponding collinear genes in *P. trichocarpa* and *R. communis*, both belong to Malpighiales, the sister group of Celastrales, indicating that these collinear genes came from the common ancestor (Figure 6). Due to *T. wilfordii* is the only species has been sequenced in Celastraceae family, we are lacking of genome information to check the other members within the family. We could speculate that *TwCYP712K2* divided from *TwCYP712K1* along the Celastraceae family speciation (<102.4 Mya) (Figure 2).

**Figure 6.**
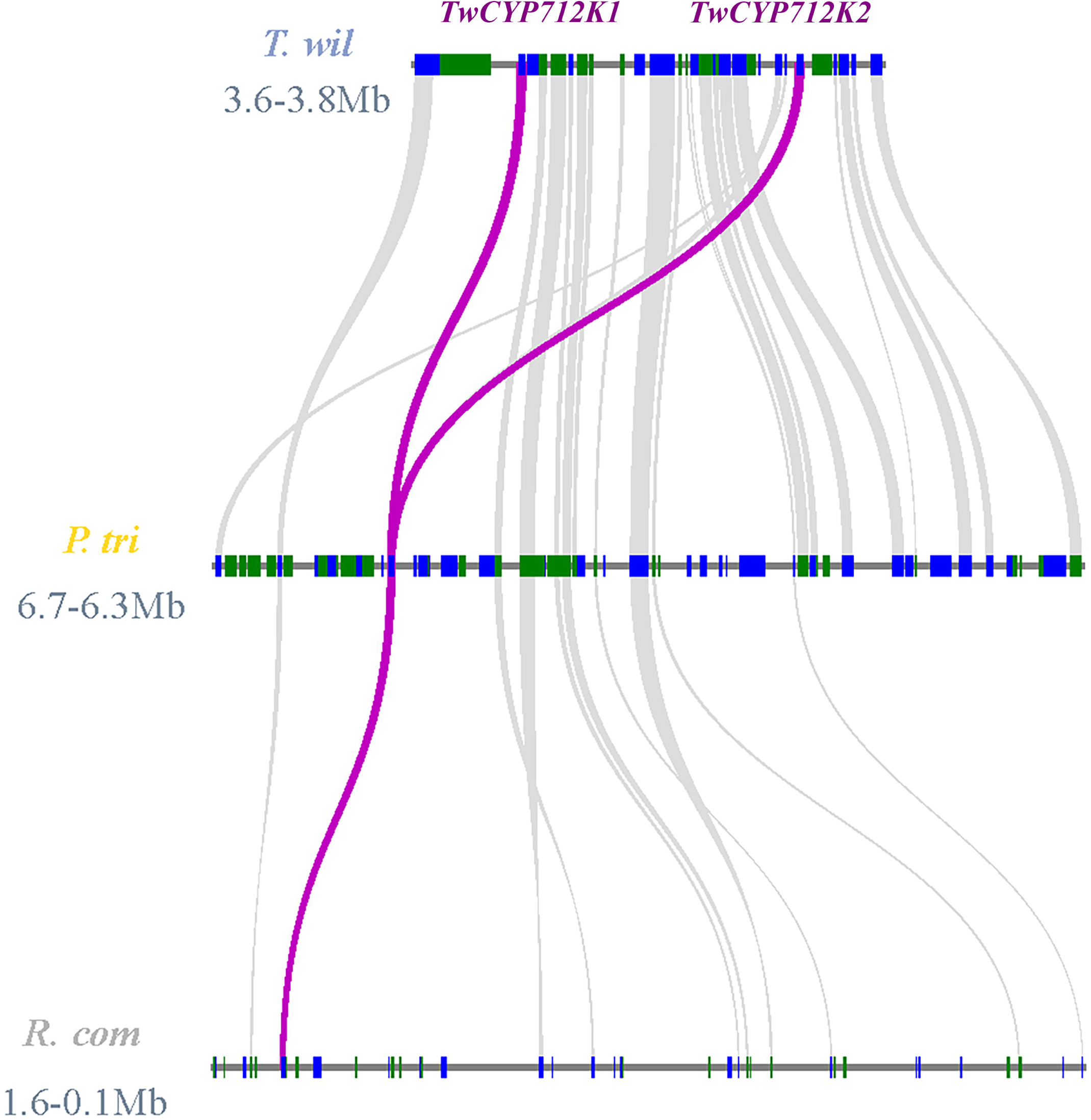
Syntenic analysis of *TwCYP712K1* and *TwCYP712K2* genes. The focused CYP genes were colored by purple. *T. wil* = *Tripterygium wilfordii*; *R. com* = *Ricinus communis*; *P. tri* = *Populus trichocarpa*.

## DISCUSSION

Friedelane-type triterpenoids are commonly distributed and richest in families Celastraceae and Hippocrateaceae (Shan et al., 2013). Celastrol is one of quinone-methide friedelanes and restricted to the Celastraceae family, which has been widely used as pharmacological ingredients in treatment of tumors, autoimmune- and obesity-related diseases (Chen et al., 2018; Ng et al., 2019; Zhao et al., 2019). However, the biosynthetic pathway of celastrol has not been fully elucidated. In this study, we provided a high-quality reference genome of *T. wilfordii* with a 340.12 Mb genome assembly and 3.09 Mb contig N50, and successfully anchored 91.02% sequences into 23 pseudochromosomes (Table 1). The quality of our genome assembly was close to the recently reported *T. wilfordii* genome (348.38 Mb total contigs and 4.36 Mb contig N50) (Tu et al., 2020).

Phylogenetic approaches and coexpression analysis have proved to be powerful tools for gene discovery in specialized metabolic pathways for nonmodel plants (Goossens, 2015; Naoumkina et al., 2010; Torrens-Spence et al., 2016). Zhou et al. (2019) isolated three OSC enzymes from *T. wilfordii*, of which TwOSC1 and TwOSC3 could convert 2, 3-oxidosqualene into friedelin as a major product, as well as β-amyrin and α-amyrin as minor products. Meanwhile, TwOSC2 catalyzed 2, 3-oxidosqualene into β-amyrin, belonging to oleanane-type triterpenoid which also found in *T. wilfordii* (Lange et al., 2017; Lv et al., 2019). However, the complete biosynthetic pathway of celastrol is still covered. In current study, based on genomic data, 35 CYP genes related to triterpenoid structure modification were identified according to the phylogenetic analysis (Figure 3), and 16 of them coexpressed with *TwOSC1* or *TwOSC3* on account of tissue-specific transcript profiles (Supplementary Figure 5 and 6). These genes could be divided into two groups that *TwOSC1* group highly expressed in leaves or other aerial parts and *TwOSC3* group specifically expressed in roots (Figure 4B), suggesting that two independent pathways may exist for friedelane-type triterpenoids biosynthesis mediated by TwOSC1 and TwOSC3, respectively. Correlation coefficients test revealed that the expression values of six *CYPs* were significantly correlated to the expression patterns of genes involved in celastrol biosynthesis and the accumulation patterns of celastrol and its biosynthetic intermediates (Figure 4C). A more subdivided phylogenetic tree showed that three putative *CYPs* were clustered closely to *CYP712K4* which was cloned from *M. ilicifolia* (Figure 4D), another plant belonged to family Celastraceae, encoding enzyme to catalyze C-29 position oxidation of friedelin to produce polpunonic acid (Bicalho et al., 2019).

*In vitro* enzyme assays revealed that TwCYP712K1 and TwCYP712K2 could use friedelin or 29-hydroxyfriedelan-3-one as substrate to produce polpunonic acid as the only product (Figure 5), indicating a two-step oxidation of friedelin at C-29 position catalyzed by the CYPs. However, we could not detect the peak of 29-hydroxyfriedelan-3-one when supplemented with friedelin, probably due to the high conversion rate between 29-hydroxyfriedelan-3-one and polpunonic acid, in consistent with the very low accumulation of 29-hydroxyfriedelan-3-one in *T. wilfordii* root (Supplementary Figure 7). Comparative genome analysis showed that *TwCYP712K1* and *TwCYP712K2* derived from the common ancestor (Figure 6). Despite they catalyzed the same reaction and located closely on the same chromosome, the identity of amino acid sequence (70.57%) was not high (Supplementary Figure 8), which suggested *TwCYP712K1* and *TwCYP712K2* were not came from recent gene duplication but separated along with the evolution of family Celastraceae. It has attracted our attentions that both of them preserved the important catalytic activity for polpunonic acid biosynthesis in *T. wilfordii*. With more genomes will be released in family Celastraceae, the evolutionary details of *TwCYP712K1* and *TwCYP712K2* can be investigated. There are many reports in which the genes encoding certain natural product pathways have been found to be grouped together in gene clusters catalyzing the biosynthesis of plant specialized metabolism including triterpenoid (Mugford et al., 2013; Nutzmann et al., 2016; Shang et al., 2014). However, neither CYPs we identified nor the signature enzyme, TwOSC1 (tw21g04301.1) and TwOSC3 (tw20g03871.1), are clustered.

In summary, we proved the integration of genome, transcriptome and metabolite analyses to be a useful strategy in screening the genes involved in plant specialized metabolism. Two CYP genes were identified catalyzing the C-29 position oxidation of friedelin to produce polpunonic acid, the biosynthetic intermediate of celastrol, which provided candidate genes for celastrol efficient production via synthetic biology.

## MATERIALS AND METHODS

### Plant materials

*T. wilfordii* plants were collected from the experimental fields of Shanghai Chenshan Botanical Garden and cultured in greenhouse by cuttage propagation. All materials used for genome sequencing were come from a single plant. For RNA-seq, roots, stems, young leaves, mature leaves, flower buds and flowers were harvested and each tissue was collected with three independent biological replicates.

### Genome survey

The features of *T. wilfordii* genome were evaluated by Illumina sequencing. Genomic DNA was isolation from leaves by DNAsecure Plant Kit (TIANGEN, China). Qualified DNA were interrupted into fragments and constructed to library using Next Ultra DNA Library Prep Kit (NEB, USA), then sequenced on the Illumina plantform. After removing adaptor reads, unidentified nucleotides (N) and low quality reads from raw reads, we estimated the genome size by performing Kmer=17 frequency analysis.

### Genome sequencing and assembly

The genome sequencing of *T. wilfordii* was performed using Oxford Nanopore Technology (ONT). Genomic DNA was extracted from leaves by modified Phenol-Chloroform method (Green and Sambrook, 2012). Qualified DNA was interrupted into fragments in g-TUBE (Covaris, USA) by a centrifuge and quality controlled by Agilent Bioanalyzer 2100. The fragmented DNA were repaired, end-prep, adapters ligated and purified using Next Ultra II End Repair/dA-Tailing Module, Next FFPE DNA Repair Mix, Next Quick Ligation Module (NEB, USA) and Agencourt AMPure XP beads (Beckman Coulter, USA), respectively. Then the qualified DNA library was sequenced on the PromethION platform.

The *de novo* assembly of genome was carried out by NextDenovo v2.1 (https://github.com/Nextomics/NextDenovo). Error correction was conducted by Racon v1.3.1 (https://github.com/isovic/racon) and Pilon v1.22 (https://github.com/broadinstitute/pilon). The completeness of the genome assembly was assessed by using BUSCO (Simão et al., 2015) and CEGMA (Parra et al., 2007), while the accuracy of assembly was evaluated by using Burrows-Wheeler Aligner (BWA) software (Li and Durbin, 2010) to align Illumina reads back to genome. Variant calling was performed using SAMtools (Li, 2011).

For Bionano sequencing, genomic DNA (molecules >300 kb) of leaves from living *T. wilfordii* plant was extracted using Plant DNA Isolation Kit (Bionano Genomics, USA). Using the NLRS DNA Labeling Kit (Bionano Genomics, USA), DNA molecules are digested with Nt.BspQI endonucleases (determined after evaluation by electronic digestion), and fluorescent labeling is performed. The labeled DNA molecules were electrophoretically stretched into linearization by Saphyr Chip (Bionano Genomics, USA) and passed through the NanoChannels, then captured on the Saphyr plantform with high-resolution camera. Raw image data was firstly converted into digital representations of the motif-specific label pattern and then analyzed using Bionano Solve v3.1 (https://bionanogenomics.com/support/software-downloads/) and its in-house scripts. Bionano data was compared to the draft genome (Nanopore version) by the following parameters: -U, -d, -T, 3, -j, 3, -N, 20, -I, 3, and scaffolds have been generated via connecting contigs when supported by the following parameters: -f, -B, 1, -N, 1.

### Sequence anchoring

Hi-C library preparation and sequencing were based on the protocol described previously with some modifications (Belton et al., 2012; Dekker et al., 2002). Leaves from living *T. wilfordii* plant were treated with 1% formaldehyde solution for fixing chromatin. Approximately 2 g fixed tissue was homogenized with liquid nitrogen and re-suspended in nuclei isolation buffer and filtered with a 40 nm cell strainer. The chromatin extraction was cut with HindIII restriction enzyme (NEB, USA), filled ends and then labeled with biotin. After ligation with T4 DNA ligase (NEB, USA) and reversing cross-linking by proteinase K, DNA was purified, interrupted into 350 bp and end-repaired. DNA fragments labeled by biotin were separated on Dynabeads M-280 Streptavidin (ThermoFisher, USA), then purified, end-repaired, A-tails added, adapter ligated and amplified by PCR to generated Hi-C libraries. At last, qualified libraries were sequenced on Illumina platform. After removing adaptor reads, unidentified nucleotides (N) and low quality reads from raw reads, the clean data was first mapped to the draft genome using BWA software (Li and Durbin, 2010). After removal of PCR duplicates and unmapped reads using SAMtools (Li et al., 2009), based on the numbers of interacted reads pairs, contigs were clustered and ordered into chromosome groups, using LACHESIS (Burton et al., 2013).

### Transcriptome sequencing

Total RNA was extracted from collected tissues using the RNAprep pure plant kit (TIANGEN, China). Qualified RNA per sample was used to generate sequencing libraries by NEBNext Ultr RNA Library Prep Kit for Illumina (NEB, USA) following manufacturer’s instruction. mRNA was purified from total RNA using poly-T oligo-attached magnetic beads, then interrupted into short fragments which were used as templates for cDNA synthesis. After purification, reparation, adenylation and adapter ligation of 3’end, 150–200 bp of cDNA fragments were separated for PCR amplification. At last, the libraries were quality control by Agilent 2100 Bioanalyzer and Q-PCR, which were then sequenced on Illumina platform. Raw reads with adapter, poly-N and low quality reads were removed to generated clean data. We mapped the RNA-seq data back to the genome assembly of *T. wilfordii* by using Bowtie v2.2.3 (Langmead and Salzberg, 2012) and TopHat v2.0.12 (Trapnell et al., 2009). The reads numbers of each gene were counted by using HTSeq v0.6.1 (Anders et al., 2014) and Fragments Per Kilobase of transcript per Million fragments mapped (FPKM) of each gene was calculated based on the length of the gene and reads count mapped to this gene (Trapnell et al., 2010).

For full-length transcriptome sequencing of PacBio, the best quality RNA samples of each tissue were mixed together to build Isoform Sequencing library by using Clontech SMARTer PCR cDNA Synthesis Kit and the BluePippin Size Selection System protocol as described by Pacific Biosciences (PN 100-092-800-03), and then sequenced on PacBio Sequel platform. Sequence data were processed using the SMRTlink 7.0 software (https://www.pacb.com/). Error correction was achieved using the Illumina RNA-seq data with the software LoRDEC (Salmela and Rivals, 2014), and redundancy in corrected consensus reads was removed by CD-HIT (Li and Godzik, 2006) to obtain final transcripts for the subsequent analysis.

### Genome annotation

Homology alignment and *de novo* prediction were applied in the repeat annotation. For homolog alignment, Repbase database employing RepeatMasker software and its in-house scripts (RepeatProteinMask) with default parameters was used to extracted repeat sequences (Jurka et al., 2005). For *de novo* prediction, LTR_FINDER (Xu and Wang, 2007), RepeatScout (Price et al., 2005), RepeatModeler (http://www.repeatmasker.org/RepeatModeler.html) with default parameters was used to build *de novo* repetitive elements database for repeat identification. Tandem repeat was also extracted by *de novo* prediction using TRF (Benson, 1999).

A combined strategy based on homology, gene prediction, RNA-seq and PacBio data were used to annotate gene structure. For homolog prediction, sequences of proteins from six species, including *A. thaliana, V. vinifera, M. truncatula, C. sativuswere, R. communis*, and *G. uralensis*, were downloaded from Ensembl/NCBI/DDBJ. Protein sequences were aligned to the genome using TblastN v2.2.26 (http://blast.ncbi.nlm.nih.gov/Blast.cgi) (E-value ≤ 1e^-5^), and then the matching proteins were aligned to the homologous genome sequences for accurate spliced alignments with GeneWise v2.4.1 (Birney et al., 2004). *De novo* gene structure identification was based on Augustus v3.2.3 (Stanke et al., 2006), GlimmerHMM v3.04 (Majoros et al., 2004), and SNAP (2013-11-29) (Korf, 2004). Based on the above prediction results, RNA-Seq reads from different tissues and PacBio reads were aligned to genome using HISAT v2.0.4 (Kim et al., 2015) and TopHat v2.0.12 (Trapnell et al., 2009) with default parameters to identify exons region and splice positions. The alignment results were then used as input for Stringtie v1.3.3 (Pertea et al., 2015) with default parameters for genome-based transcript assembly. The alignment results were then integrated into a non-redundant gene set using EVidenceModeler v1.1.1, then further corrected with PASA to predict untranslated regions and alternative splicing to generate the final gene set (Haas et al., 2008).

According to the final gene set, gene function was predicted by aligning the protein sequences to the SwissProt (http://www.uniprot.org/), Nr (http://www.ncbi.nlm.nih.gov/protein), Pfam (http://pfam.xfam.org/), KEGG (http://www.genome.jp/kegg/), and InterPro v5.31 (https://www.ebi.ac.uk/interpro/) using Blastp (E-value ≤ 1e^-5^). The Gene Ontology (GO) IDs for each gene were assigned according to the corresponding InterPro entry.

Non-coding RNA was annotated using tRNAscan-SE (for tRNA) (Lowe and Chan, 2016) or INFERNAL (for miRNA and snRNA) (Nawrocki and Eddy, 2013). rRNA was predicted by BLAST using rRNA sequences from *A. thaliana* and *O. sativa* as references, which were highly conserved among plants.

### Comparative genome analyses

Gene families clustering of 12 sepecies, including *Tripterygium wilfordii, Arabidopsis thaliana, Citrus sinensis, Vitis vinifera, Glycine max, Medicago truncatula, Glycyrrhiza uralensis, Cucumis sativus, Populus trichocarpa, Ricinus communis, Oryza sativa*, and *Amborella trichopoda* were inferred through all-against-all protein sequence similarity searches using OthoMCL (Li et al., 2003) with E-value of 1e^-5^. Proteins less than 50 amino acids were removed and were retained only the longest predicted transcript per locus. All the proteins were clustered by OrthoMCL with inflation 1.5.

Single-copy orthologous genes were retrieved from the 12 species and aligned using MUSCLE (Edgar, 2004). All the alignments were combined together to produce a super alignment matrix, which was used to construct a maximum likelihood (ML) phylogenetic tree using RAxML (Stamatakis, 2006).

Divergence times between species were calculated using MCMCtree program implemented in the PAML (Yang, 2007). The following calibration points were applied: *M. truncatula* – *G. uralensis* (15-91 Mya), *G. max* – *M. truncatula* (46-109 Mya), *G. max* – *C*.*sativus* (95-135 Mya), *A. thaliana* – *C. sinensis* (96-104 Mya), *P. trichocarpa* – *R. communis* (70-86 Mya), *A. thaliana* – *P. trichocarpa* (98-117 Mya), *C. sativus* – *R. communis* (101-131 Mya), *V. vinifera* – *A. thaliana* (107-135 Mya), *V. vinifera* – *O. sativa* (115-308 Mya), *O. sativa* – *A. trichopoda* (173-199 Mya), which were extracted from TimeTree (http://www.timetree.org/).

Expansion and contraction of gene families were analyzed by using CAFE (Bie et al., 2006) with *P*-value threshold 0.05 and 10000 random samples. In order to avoid false positive, the results were filtered and the enrichment results were screened with Family-wide *P*-value <0.05 and Viterbi *P*-values <0.05.

### Genome-wide identification of CYP450 genes

Hidden Markov model (HMM) profile of Pfam PF06200 (http://pfam.sanger.ac.uk) was used to extract full-length CYP candidates from *T. wilfordii* genome by the HMM algorithm (HMMER) (Eddy, 1998), filtering by a length between 400 and 600 amino acids (Christ et al., 2019).

### Phylogenetic analyses

Multiple sequences alignments and phylogenetic trees construction were performed by MEGA X (Kumar et al., 2018), using either Neighbor-Joining or Maximum-likelihood methods with a bootstrap test (n=1000 replications).

### Coexpression analysis

Gene expression pattern analysis was perform by Short Time-series Expression Miner software (STEM) (Ernst and Bar-Joseph, 2006) on the OmicShare tools platform (www.omicshare.com/tools). The parameters were set as follows: maximum unit change in model profiles between time points is 1; maximum output profiles number is 20 (similar profiles will be merged); minimum ratio of fold change of DEGs is no less than 2.0 and with *P*-value<0.05.

### Gene cloning

The complete ORFs of the putative CYP genes were amplified by the primers listed in Supplemental Table 14, using cDNA from *T. wilfordii* root as template. According to the manufacturer’s instructions, the fragments were cloned into entry vector pDONR207 and yeast expression vector pYesdest52 using Gateway BP Clonase II Enzyme Kit and LR Clonase II Enzyme Kit (Invitrogen, MA, USA), respectively.

### Standard compounds

Friedelin, 29-hydroxy-friedelan-3-one and celastrol were purchased from Yuanye-biotech (Shanghai, China), polpunonic acid and wilforic acid A were purchased from Weikeqi-biotech (Sichuan, China). Friedelin was dissolved in DMSO/isopropanol (v/v=1:2) following half hour ultrasonic water bath, while 29-hydroxy-friedelan-3-one, celastrol, polpunonic acid and wilforic acid A were dissolved in methanol.

### Enzyme assays of yeast *in vivo*

Yeast *in vivo* assays was followed previously described protocol with some modifications (Zhao et al., 2018). The yeast expression vector constructs or empty vector were transformed into yeast *Saccharomyces cerevisiae* WAT11 (Pompon et al., 1996; Truan et al., 1993) by using Yeast Transformation II Kit (ZYMO, CA, USA) and screened on synthetic SD-dropout medium lacking uracil (SD-Ura) with 20 g/L glucose. After growing at 28°C for 48-72 h, transformant colonies were initially grown in 20 ml of SD-Ura liquid medium with 20 g/L glucose at 28°C for about 24 h until OD_600_ reached 2 to 3. Yeast cells were harvested by centrifugation with 4000 rpm and resuspended in 20 mL SD-Ura liquid medium supplemented with 20 g/L galactose for inducing of target proteins, while friedelin or 29-hydroxy-friedelane-3-one was applied to the cultures at a 25 mM final concentration. After 48 h fermentation (supplemented 2 mL galactose after 24 h), yeast cells were harvested by centrifugation and extracted with 2 mL 70% methanol by an ultrasonic water bath for 2 h. The supernatants were filtered with 0.2 μm millipore filter and analyzed by LC–MS.

### Enzyme assays *in vitro*

Protocol of enzyme assays *in vitro* was followed as described previously with some modifications (Zhao et al., 2016b). Yeast transformation and target proteins induction were same as described above, except for the 24 h of fermentation after galactose supplemented. Yeast cells were harvested by centrifugation and suspended with 10 mL mixture of 50 mM Tris-HCl (pH 7.5), 1 mM EDTA, 0.5 mM phenylmethylsulfonyl fluoride, 1 mM dithiothreitol, 0.6 M sorbitol and ddH_2_O. High pressure cell disruption equipment (Constant Systems, Northants, UK) was used to crush the yeast cells. After centrifugation, about 10 mL supernatants were collected and applied with CaCl_2_ at an 18 mM final concentration. Microsomal proteins were then collected by centrifugation and suspended in a storage buffer containing 50 mM Tris-HCl (pH 7.5), 1 mM EDTA and 20% (v/v) glycerol with a final concentration of 10 to 15 mg/mL measuring by the Bradford method (Bradford, 1976).

The catalytic activity of putative CYP was assayed in a 100 μl reaction volume, which contained 100 mM sodium phosphate buffer (pH 7.9), 0.5 mM reduced glutathione, 2.5 μg extracted protein and 100 μM substrate (friedelin or 29-hydroxy-friedelan-3-one). The reaction was initiated by adding NADPH at 1 mM and incubated for 12 h at 28 °C. Methanol was then added to a final concentration of 70% to quench the reaction. The reaction mixture was filtered with 0.2 μm millipore filter and analyzed by LC–MS. Microsomal proteins extracted from yeast harboring the empty vector was set as a negative control.

### Metabolite analysis

Plant tissue was ground into powder in liquid nitrogen and then freeze dried. Fifty mg sample was suspended in 2 ml of 80% (v/v) methanol, setting overnight at room temperature, and then extracted by an ultrasonic water bath for 60 min. After 12000 rpm centrifugation for 2 min, the supernatant was filtered through a 0.2 μm millipore filter before LC-MS analysis.

The contents of celastrol and wilforic acid A were analyzed by Agilent 1260LC-6400 QQQ (triple quadrupole mass). The chromatographic separation was carried out on an Agilent Eclipse XDB-C18 analytical column (4.6×250 mm, 5μm) with a guard-column. The mobile phase consisting of of 0.1% (v/v) formic acid in water (A) and acetonitrile (B) was set at a flow rate of 0.8 mL/min. The gradient program was as follows: 0-12 min, 10-60% B; 12-17 min, 70% B; 17-25 min, 95% B; 25-28 min, 95% B; 28-29 min, 5% B; 29-35 min, 5% B. The detection wavelength of celastrol was 425 nm, and UV spectra from 190 to 500 nm were also recorded. The injection volume was 10 μl and column temperature was 35 °C. The LC effluent was introduced into the ESI source by a splite-flow valve with ratio of 3:1. All mass spectra were acquired in the negative ion mode, and the parameters were as follows: dry gas 4 L/min; dry gas temperature 300 °C; Nebulizer (high-purity nitrogen) pressure 15 psi; capillary voltage 4.0 kV; fragmentor 135; cell accelerator voltage 7. For full-scan MS analysis, the spectra were recorded in the range of m/z 100–750.

The contents of 29-hydroxy-friedelan-3-one and polpunonic acid were analyzed by Thermo Q Exactive Plus. The chromatographic separation was carried out on a Thermo Syncronis C18 column (2.1 × 100 mm, 1.7μm). The mobile phase consisting of 0.1% (v/v) formic acid in water (A) and acetonitrile (B) was set at a flow rate of 0.4 mL/min. The gradient program was as follows: 0-12 min, 10-60% B; 12-17 min, 70% B; 17-25 min, 95% B; 25-28 min, 95% B; 28-29 min, 5% B. Mass spectra were acquired in both positive and negative ion mode with HESI source, and the parameters were as follows: aus. gas flow 10 L/min; aus. gas heater 350°C; sheath gas flow 40 L/min; spray voltage 3.5 kV; capillary temperature 320°C. For full-scan MS/ddMS^2^ analysis, the spectra were recorded in the range of m/z 50–750 at a resolution of 17500 with AGC target 1e^6^ and 2e^5^, respectively.

The contents of metabolites in different tissues were measured by comparing the area of the individual peaks with standard curves obtained from standard compounds.

### Syntenic analyses

The genomes of *T. wilfordii, P. trichocarpa* and *R. communis* were compared with MCScan Toolkit v1.1 (Wang et al., 2012) implemented in python. Genomes of *P. trichocarpa* v4.1 and *R. communis* v0.1 were downloaded from phytozome v13 (https://phytozome-next.jgi.doe.gov/). Syntenic gene pairs were identified using all-vs-all BLAST search using LAST (Frith et al., 2010), filtered to remove pairs with scores below 0.7, and clustered into syntenic blocks in MCScan. Microsynteny plots were constructed using MCScan.

## DATA AVAILABILITY

Reference genome and RNA-seq data are deposited in GenBank under project number PRJNA640746. Gene and protein sequences of TwCYP712K1 (MT633088) and TwCYP712K2 (MT633089) are deposited in GenBank.

## COMPLIANCE AND ETHICS

The authors declare that they have no conflict of interest.

## ACKNOWLEDGEMENTS

This work was supported by the National Key R&D Program of China (2018YFC1706202, 2019YFD1000703, 2018YFD1000701), the National Natural Science Foundation of China (31870282, 31700268), Youth Innovation Promotion Association CAS, and the Chenshan Special Fund for Shanghai Landscaping Administration Bureau Program (G182401, G182402, G192419, G192413, G192414 and G202402). Qing is also support by The Shanghai Youth Talent Support Program and SA-SIBS Scholarship Program. We greatly appreciate facilities and services office of Chenshan Plant Science Research Center for experimental instruments supporting.

