## Supplemental Figures&Tables for "The genome analysis of *Tripterygium wilfordii* reveals TwCYP712K1 and *TwCYP712K2* responsible for oxidation of friedelin in celastrol biosynthesis pathway"

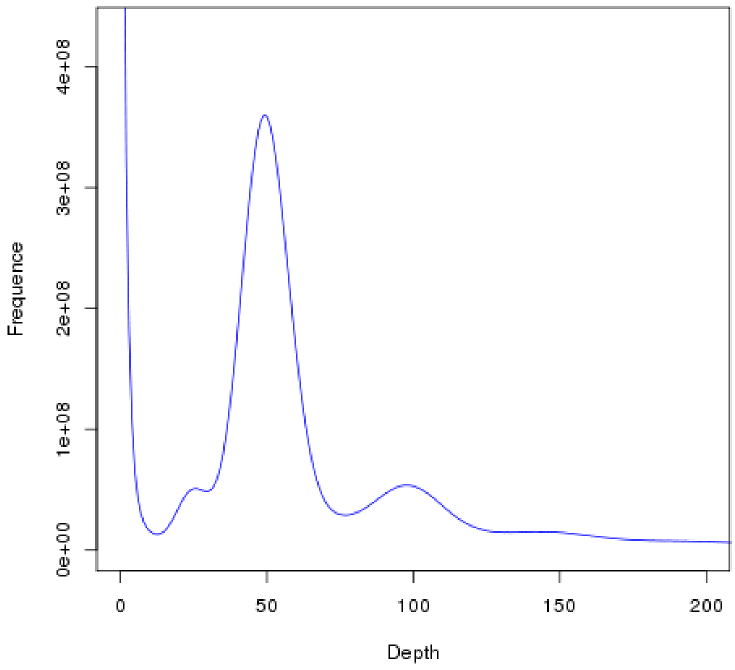


**Supplementary Figure 1. Estimation of *T. wilfordii* genome size by k-mer analysis**

X axis shows k-mer depth and Y axis shows k-mer frequency. G0=k-mer number/depth, G=G0*(1-Error rate) where G0 is previous genome size and G is revised genome size. The genome size was measured as 375.84 Mb using this method.

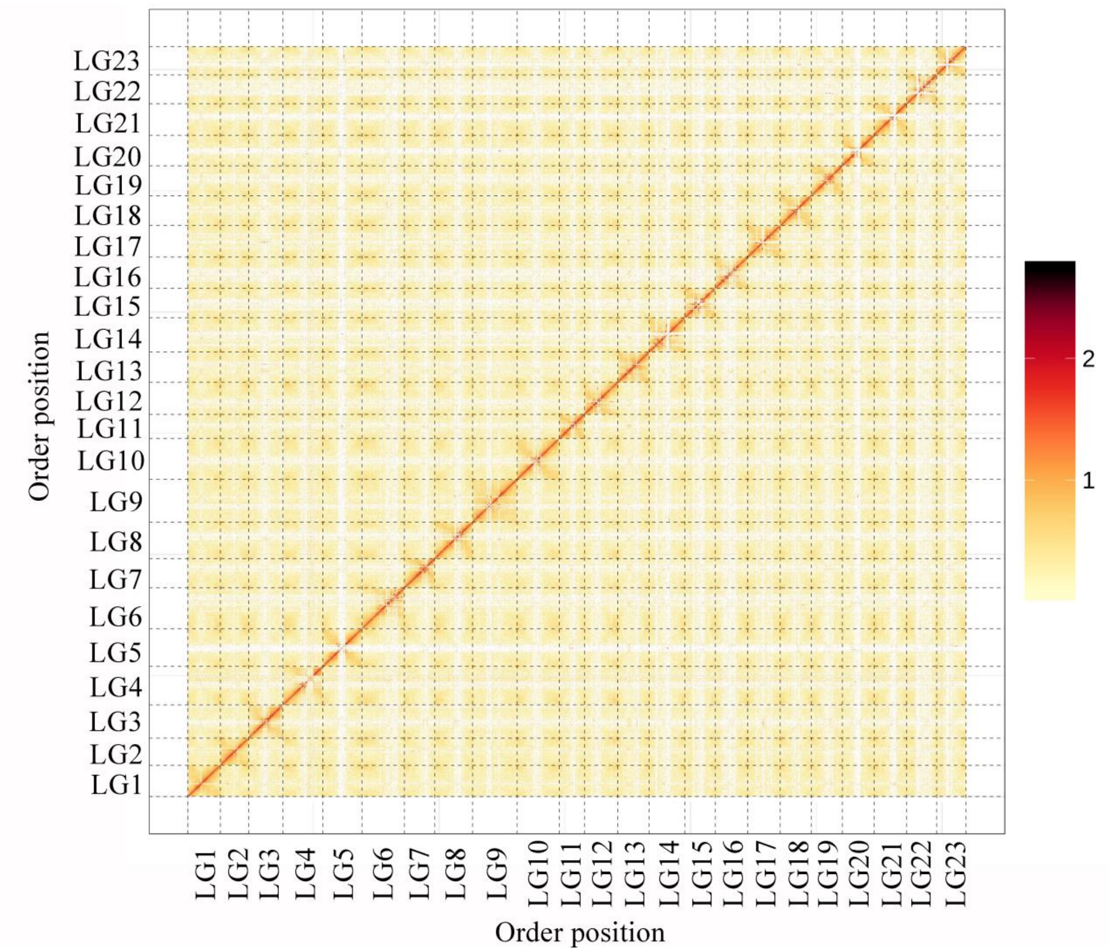


**Supplementary Figure 2. Interaction heat-map of chromosomal fragments based on Hi-C analysis**

LG1-LG23 indicate Lachesis Groups 1-23. X and Y axes indicate the order positions of scaffolds on corresponding pseudochromosomes. The bar represents interaction strength between sequence segments.

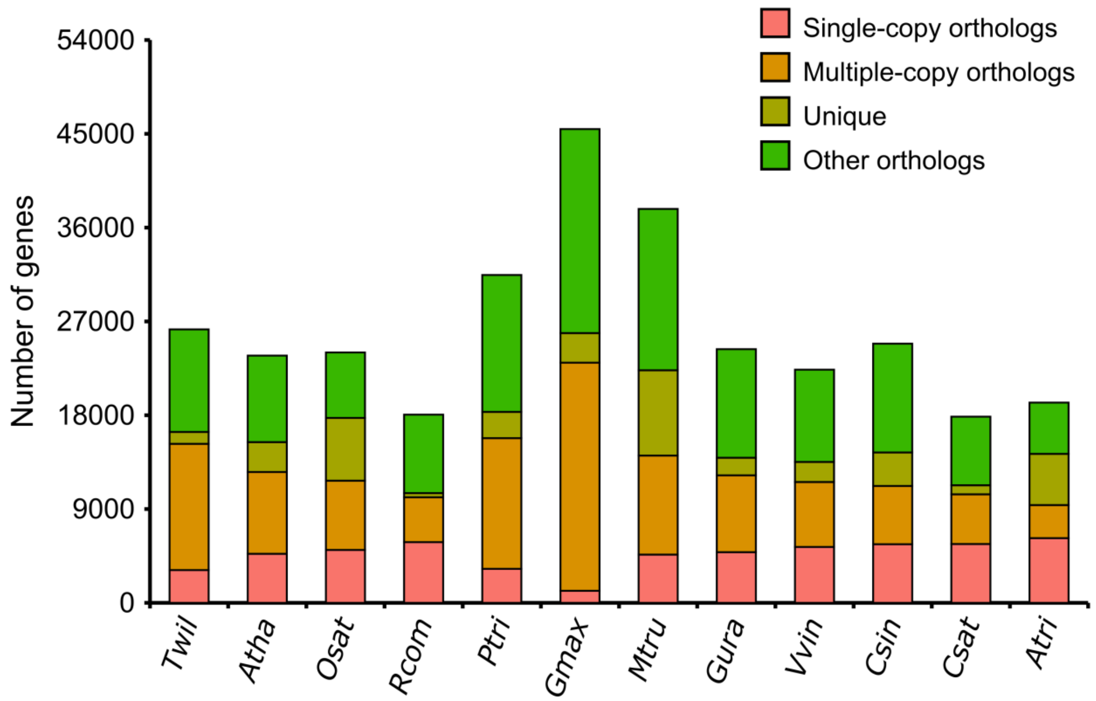


**Supplementary Figure 3. The distribution of genes in different species**

*Twil* = *Tripterygium wilfordii*; *Atha* = *Arabidopsis thaliana*; *Osat* = *Oryza sativa*; *Rcom* = *Ricinus communis*; *Ptri* = *Populus trichocarpa*; *Gmax* = *Glycine max*; *Mtru* = *Medicago truncatula*; *Gura*= *Glycyrrhiza uralensis*; *Vvin* = *Vitis vinifera*; *Csin* = *Citrus sinensis*; *Csat* = *Cucumis sativus*; *Atri* = *Amborella trichopoda*.

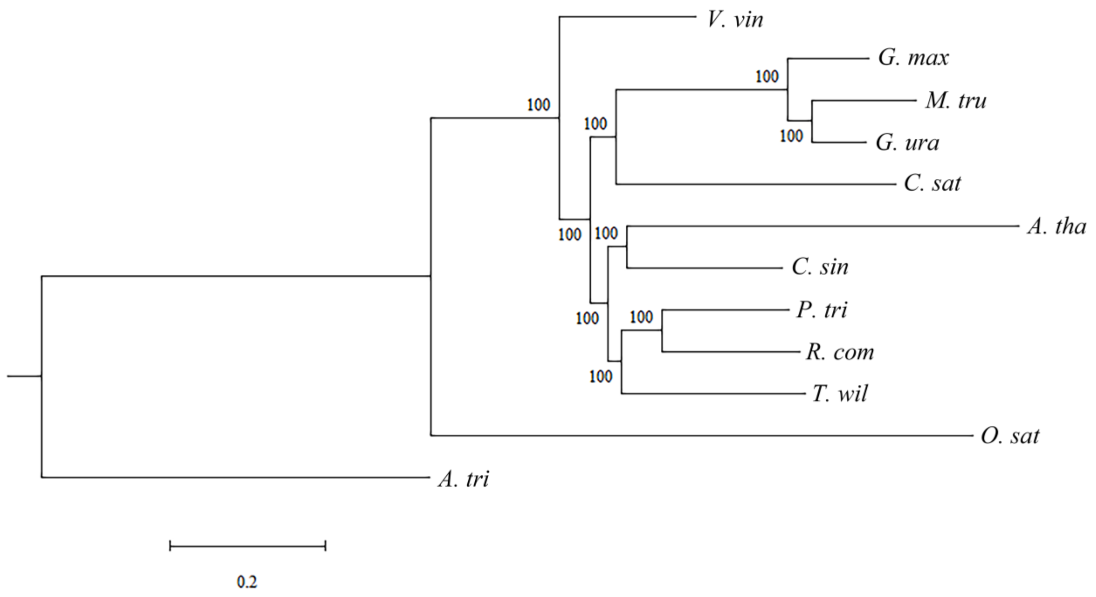


**Supplementary Figure 4. Phylogenetic tree of *T. wilfordii* and other selected species.**

The branch length represents the evolution rate, and the value on the branch represents the value of bootstrap support. *T. wil* = *Tripterygium wilfordii*; *A. tha* = *Arabidopsis thaliana*; *O. sat* = *Oryza sativa*; *R. com* = *Ricinus communis*; *P. tri* = *Populus trichocarpa*; *G. max* = *Glycine max*; *M. tru* = *Medicago truncatula*; *G. ura*= *Glycyrrhiza uralensis*; *V. vin* = *Vitis vinifera*; *C. sin* = *Citrus sinensis*; *C. sat* = *Cucumis sativus*; *A. tri* = *Amborella trichopoda*.

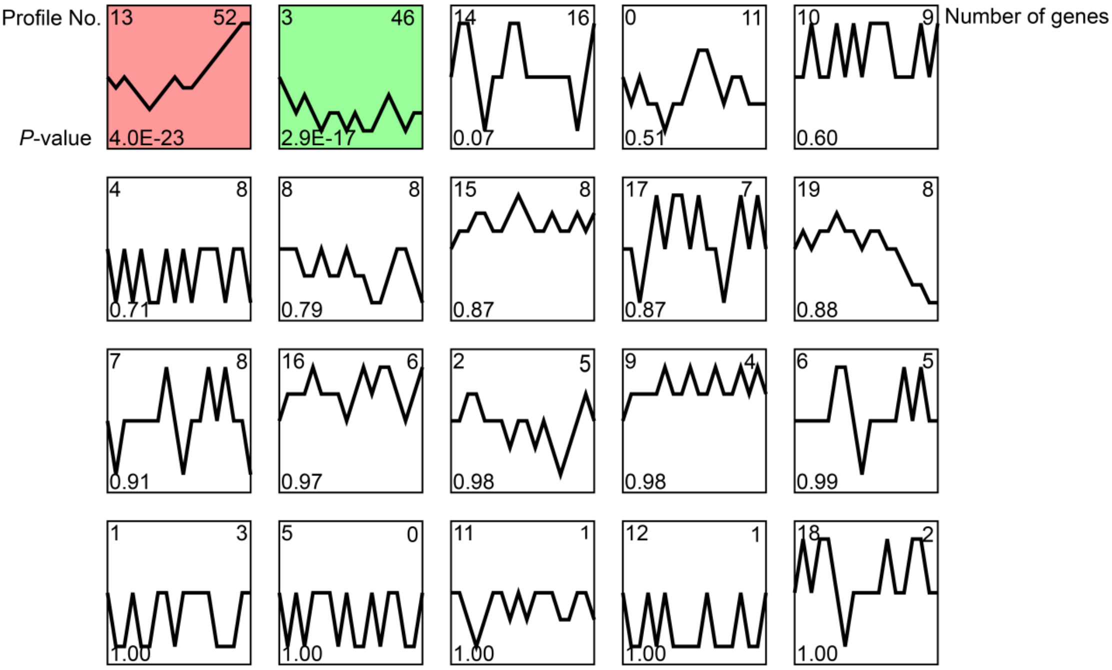


**Supplementary Figure 5. Maps of expression trends of CYPs with TwOSC1 and TwOSC3**

Profiles ordered based on the *P*-value significance of number of genes assigned versus expected. Numbers on the top left corner represent the profiles number; numbers on the left bottom represent the *P*-value; numbers on the top right corner represent the total number of genes. Colored maps represent the significant enrichment with *P*-value<0.05.

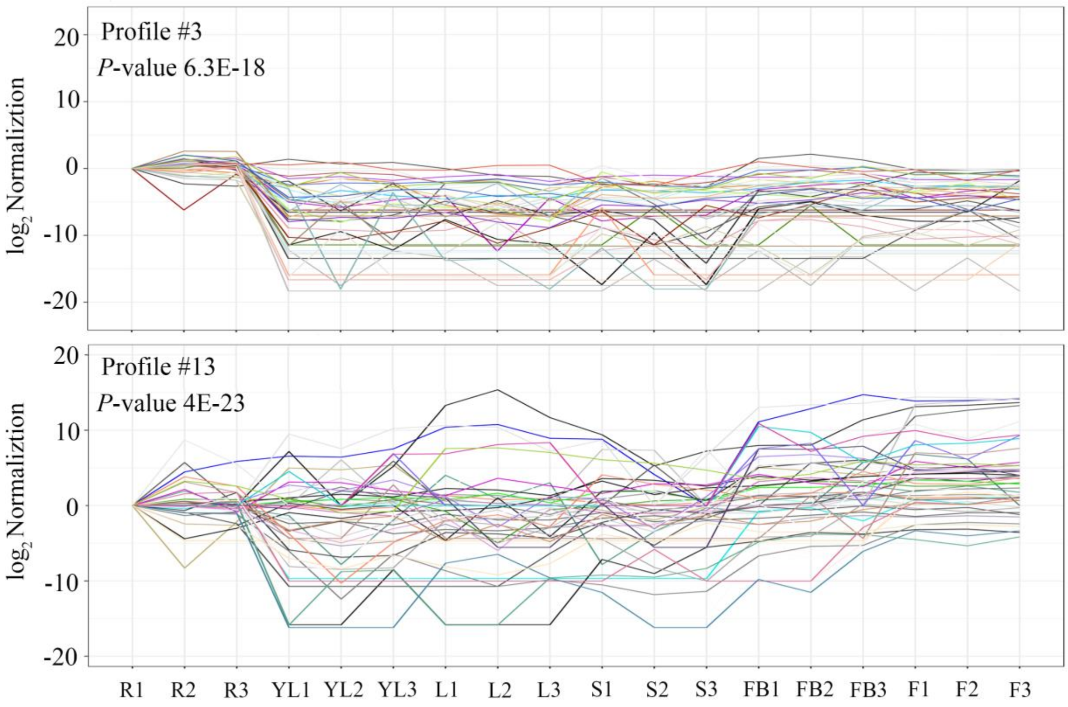


**Supplementary Figure 6. Coexpression of potential CYP genes with *TwOSC1* and *TwOSC3***

The upper map indicates the similar expression patterns of *CYPs* with *TwOSC3* and the lower map indicates the similar expression patterns of *CYPs* with *TwOSC1*. R, root; YL, young leaf; L, leaf; S, stem; FB, flower bud; F, flower; numbers 1-3 represents three biological replicates, respectively.

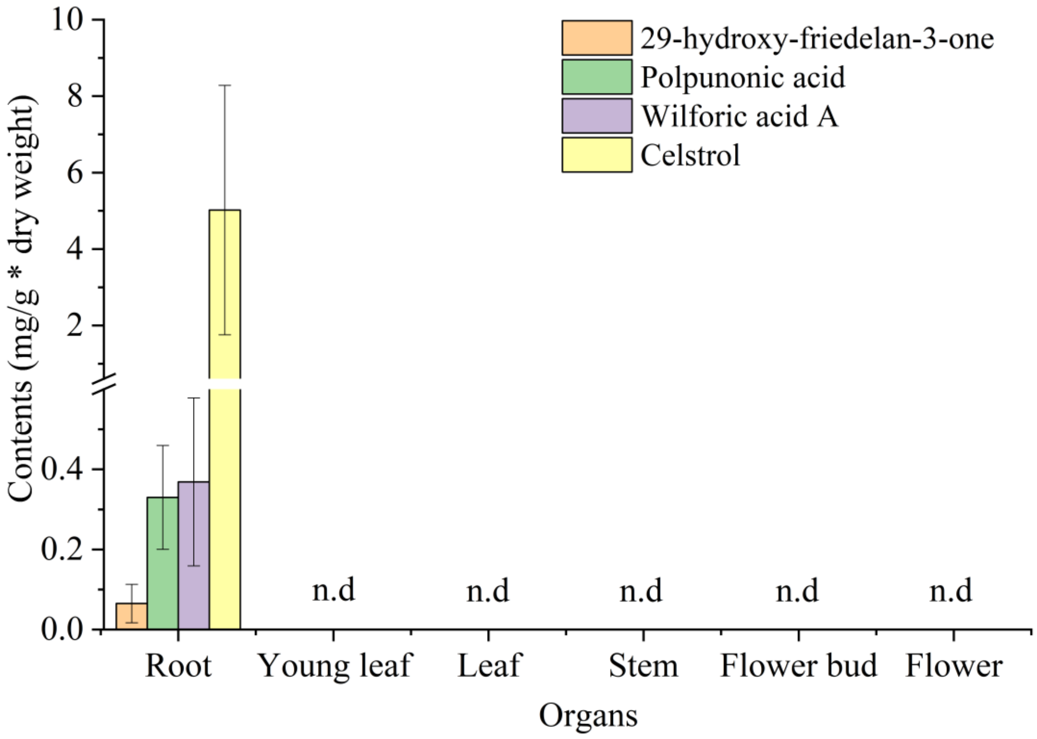


**Supplementary Figure 7. The accumulation of celastrol and intermediate products in different tissues of *T. wilfordii***

Bars are means ± SD from three independent biological replicates, n.d=not detected with our experimental condition, and the value of not detected compounds were set to 0.

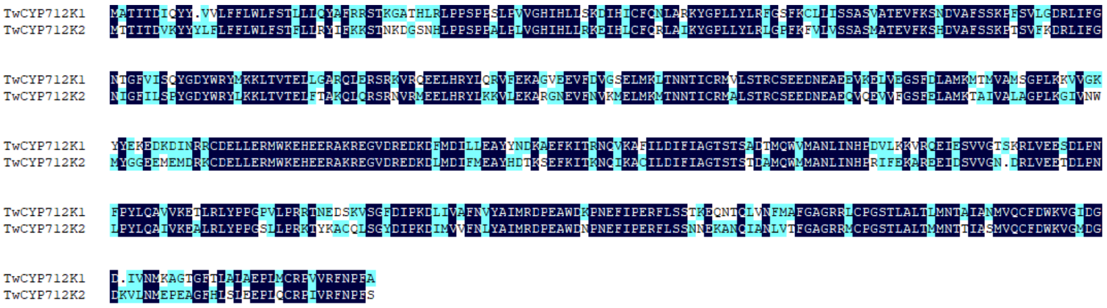

**Supplementary Figure 8. Alignment of TwCYP712K1 and TwCYP712K2 sequences**

TwCYP712K1 and TwCYP712K2 exhibited 70.57% identity. The consensus sequences were highlighted by deep blue color.

**Supplementary Table 1. Genome sequencing data and sequencing coverage**

| Pair-end libraries | Total data (Gb) | Sequence coverage (X) |
|---|---|---|
| Illumina Hiseq PE150 (for genome survey and error correction) | 25.32 | 67.37 |
| Nanopore | 77.86 | 207.16 |
| BioNano | 60.80 | 161.77 |
| Illumina Hiseq PE150 (for Hi-C) | 77.68 | 206.68 |
| PacBio (for annotation) | 20.75 | 55.21 |

**Supplementary Table 2. Statistics of genome assembly**

| Sample ID | Nanopore version | | | | Bionano hybrid scaffold version | | | |
|---|---|---|---|---|---|---|---|---|
| | length | | number | | length | | number | |
| | Contig* (bp) | Scaffold (bp) | Contig* | Scaffold | Contig* (bp) | Scaffold (bp) | Contig* | Scaffold |
| Total | 340,124,188 | - | 553 | - | 340,124,188 | 342,588,429 | 553 | 470 |
| Max | 7,962,777 | - | - | - | 7,962,777 | 10,510,391 | - | - |
| Number>=2000 | - | - | 553 | - | - | - | 553 | 470 |
| N50 | 3,088,446 | - | 34 | - | 3,088,446 | 5,425,714 | 34 | 25 |
| N60 | 2,351,287 | - | 46 | - | 2,351,287 | 4,027,572 | 46 | 32 |
| N70 | 1,048,940 | - | 68 | - | 1,048,940 | 3,189,075 | 68 | 41 |
| N80 | 334,278 | - | 133 | - | 334,278 | 634,293 | 133 | 63 |
| N90 | 202,837 | - | 264 | - | 202,837 | 205,847 | 264 | 179 |

* Contigs after scaffolding

**Supplementary Table 3. Nucleotide statistics in the draft genome assembly**

| | Number (bp) | % of genome |
|---|---|---|
| A | 106,752,681 | 31.39 |
| T | 106,872,067 | 31.42 |
| C | 63,093,649 | 18.55 |
| G | 63,405,791 | 18.64 |
| N | 0 | 0.00 |
| Total | 340,124,188 | - |
| GC | 126,499,440 | 37.19 |

**Supplementary Table 4. SNP statistics of the genome assembly**

| | Number | Percentage |
|---|---|---|
| All SNP | 766560 | 0.256773% |
| Heterozygosis SNP | 756672 | 0.253461% |
| Homology SNP | 9888 | 0.003312% |

**Supplementary Table 5. Summary of BUSCO evaluation**

| | Percentage (%) |
|---|---|
| Complete BUSCOs | 95.2 |
| Complete and single-copy BUSCOs | 77.7 |
| Complete and duplicated BUSCOs | 17.5 |
| Fragmented BUSCOs | 1.1 |
| Missing BUSCOs | 3.7 |
| Total BUSCO groups searched | 1440 |

**Supplementary Table 6. Summary of CEGMA evaluation**

| | Complete | | Complete + Partial | |
|---|---|---|---|---|
| **Species** | # Prots | %Completeness | # Prots | %Completeness |
| *T. wilfordii* | 233 | 93.95 | 238 | 95.97 |

Complete: core gene >70% assembly; Complete + partial: core gene partially assembly; #Prots: numbers of core gene assembly; %completeness: ration of assembled core gene to core gene library.

**Supplementary Table 7. Summary of short reads coverage of genome assembly**

| | | % of Percentage |
|---|---|---|
| Reads | Mapping rate (%) | 95.31 |
| | Average sequencing depth | 63.05 |
| | Coverage (%) | 93.99 |
| Genome | Coverage at least 4X (%) | 92.32 |
| | Coverage at least 10X (%) | 90.59 |
| | Coverage at least 20X (%) | 87.59 |

**Supplementary Table 8. Statistics of Hi-C assembly**

| Sample ID | Contig length | Scaffold length | Contig number* | Scaffold number |
|---|---|---|---|---|
| Total | 340,124,188 | 342,608,929 | 566 | 279 |
| Max | 7,962,777 | 17,748,360 | - | - |
| Number>=2000 | - | - | 566 | 279 |
| N50 | 2,929,360 | 13,028,512 | 36 | 12 |
| N60 | 2,234,477 | 12,513,880 | 49 | 15 |
| N70 | 926,087 | 12,436,525 | 73 | 17 |
| N80 | 332,138 | 11,980,432 | 140 | 20 |
| N90 | 202,098 | 10,152,376 | 271 | 23 |

*Contigs>100 bp were selected for statistics

**Supplementary table 9. Scaffold number and length grouped on pseudochromosomes**

| Group | Number of scaffold | Total Length (bp) |
|---|---|---|
| Group1 | 2 | 13,014,687 |
| Group2 | 4 | 11,285,713 |
| Group3 | 9 | 13,837,602 |
| Group4 | 19 | 16,052,000 |
| Group5 | 7 | 15,585,988 |
| Group6 | 12 | 16,997,832 |
| Group7 | 4 | 12,198,226 |
| Group8 | 11 | 15,126,992 |
| Group9 | 8 | 17,748,360 |
| Group10 | 6 | 16,856,595 |
| Group11 | 9 | 10,152,376 |
| Group12 | 8 | 13,417,622 |
| Group13 | 12 | 12,513,880 |
| Group14 | 17 | 14,247,452 |
| Group15 | 12 | 12,206,946 |
| Group16 | 14 | 13,028,512 |
| Group17 | 12 | 13,091,630 |
| Group18 | 5 | 12,436,525 |
| Group19 | 12 | 12,453,453 |
| Group20 | 9 | 12,746,742 |
| Group21 | 8 | 13,078,104 |
| Group22 | 15 | 11,980,432 |
| Group23 | 13 | 11,790,904 |
| Total | 228 | 311,848,573 (91.02%) |

**Supplementary Table 10. Summary of gene structure annotation**

| | Gene set | Number | Average transcript length (bp) | Average CDS length (bp) | Average exons per gene | Average exon length (bp) | Average intron length (bp) |
|---|---|---|---|---|---|---|---|
| *De novo* | Augustus | 28,686 | 3,220.73 | 1,220.87 | 5.18 | 235.80 | 478.71 |
| | GlimmerHMM | 48,445 | 5,104.71 | 779.48 | 3.35 | 233.02 | 1,844.39 |
| | SNAP | 38,054 | 2,620.07 | 821.85 | 4.29 | 191.65 | 546.86 |
| | Geneid | 41,284 | 4,097.53 | 941.16 | 4.72 | 199.31 | 848.01 |
| | Genscan | 27,524 | 7,575.88 | 1,391.54 | 6.72 | 207.03 | 1,080.88 |
| Homolog | Rco | 25,778 | 2,920.77 | 1,170.30 | 5.07 | 230.82 | 430.08 |
| | Gur | 24,898 | 2,858.12 | 1,121.45 | 4.87 | 230.21 | 448.59 |
| | Vvi | 25,299 | 2,873.47 | 1,140.76 | 5.03 | 226.77 | 429.90 |
| | Ath | 24,236 | 2,814.33 | 1,131.59 | 4.94 | 229.01 | 426.97 |
| | Csa | 25,267 | 2,748.49 | 1,125.37 | 4.86 | 231.64 | 420.68 |
| | Mtr | 25,346 | 2,771.60 | 1,111.88 | 4.87 | 228.30 | 428.84 |
| RNASeq | PASA | 123,067 | 3,258.88 | 1,066.20 | 5.26 | 202.69 | 514.67 |
| | Transcripts | 48,394 | 6,624.21 | 2,133.18 | 6.71 | 317.76 | 786.09 |
| EVM | | 34,739 | 3,020.48 | 1,106.33 | 4.91 | 225.17 | 489.13 |
| Pasa-update* | | 34,427 | 3,016.92 | 1,130.24 | 4.97 | 227.63 | 475.81 |
| Final set* | | 31,593 | 3,180.62 | 1,182.78 | 5.22 | 226.73 | 473.80 |

* containing UTR region

Rco: *Ricinus communis*; Gur: *Glycyrrhiza uralensis*; Vvi: *Vitis vinifera*; Ath: *Arabidopsis thaliana*; Csa: *Cucumis sativus*; Mtr: *Medicago truncatula*

**Supplementary Table 11. Summary of gene function annotation**

| | Number | Percent(%) |
|---|---|---|
| Total | 31,593 | - |
| Swissprot | 25,392 | 80.40 |
| Nr | 30,388 | 96.20 |
| KEGG | 24,509 | 77.60 |
| InterPro | 29,587 | 93.70 |
| GO | 17,963 | 56.90 |
| Pfam | 24,502 | 77.60 |
| Annotated | 30,535 | 96.70 |
| Unannotated | 1,058 | 3.30 |

**Supplementary Table 12. Summary of repetitive sequences**

| | **Denovo+Repbase*** | | **TE Proteins**** | | **Combined TEs***** | |
|---|---|---|---|---|---|---|
| | Length(bp) | % in Genome | Length(bp) | % in Genome | Length(bp) | % in Genome |
| DNA | 5,485,526 | 1.60 | 464,373 | 0.14 | 5,755,303 | 1.68 |
| LINE | 2,270,719 | 0.66 | 161,136 | 0.05 | 2,365,709 | 0.69 |
| SINE | 661,709 | 0.19 | 0 | 0 | 661,709 | 0.19 |
| LTR | 124,801,666 | 36.43 | 24,680,784 | 7.20 | 125,876,630 | 36.74 |
| Unknown | 17,022,433 | 4.97 | 0 | 0 | 17,022,433 | 4.97 |
| Total | 147,540,202 | 43.06 | 25,305,396 | 7.39 | 148,553,910 | 43.36 |

* Denovo+Repbase: Integrated the results of RepeatModeler, RepeatScout, Piler, LTR FINDER and RepBase, then annotated by RepeatMasker; ** TE Proteins: annotated by RepeatProteinMask based on RepBase; *** Combined TEs: Integrated the results of Denovo+Repbase and TE Proteins, and then removed redundant

**Supplementary Table 13. Summary of non-coding RNA**

| Type | | Copy number | Average length (bp) | Total length (bp) | % of genome |
|---|---|---|---|---|---|
| | miRNA | 355 | 120.26 | 42,694 | 0.012461 |
| | tRNA | 797 | 75.45 | 60,134 | 0.017552 |
| | rRNA | 827 | 307.68 | 254,452 | 0.074269 |
| | 18S | 271 | 690.83 | 187,216 | 0.054644 |
| rRNA | 28S | 293 | 125.56 | 36,790 | 0.010738 |
| | 5.8S | 104 | 131.08 | 13,632 | 0.003979 |
| | 5S | 159 | 105.75 | 16,814 | 0.004908 |
| | snRNA | 982 | 109.09 | 107,128 | 0.031268 |
| | CD-box | 777 | 101.82 | 79,113 | 0.023091 |
| snRNA | HACA-box | 90 | 129.56 | 11,660 | 0.003403 |
| | splicing | 114 | 142.35 | 16,228 | 0.004737 |
| | scaRNA | 1 | 127 | 127 | 0.000037 |

**Supplementary Table 14. Primers used for genes cloning**

| Primer names | Squence (5' to 3') * |
|---|---|
| TwCYP712K1-F | GGGGACAAGTTTGTACAAAAAAGCAGGCTTCATGGCCACCATCACTGACATC |
| TwCYP712K1-R | GGGGACCACTTTGTACAAGAAAGCTGGGTTTTAACCGGCAAATGGATTGAA |
| TwCYP712K2-F | GGGGACAAGTTTGTACAAAAAAGCAGGCTTCATGACAACAATCACTGATGTGAA |
| TwCYP712K2-R | GGGGACCACTTTGTACAAGAAAGCTGGGTTTTAAGAAGAAAATGGATTGAACC |
| TwCYP712K3-F | GGGGACAAGTTTGTACAAAAAAGCAGGCTTCATGGCCACCACTACCATCATT |
| TwCYP712K3-F | GGGGACCACTTTGTACAAGAAAGCTGGGTTTTAGCAAGAAAAGGGATGGAATC |

* Underlined sequences represent Gateway prime
